# Rapid Evolution of SARS-CoV-2 Challenges Human Defenses

**DOI:** 10.1101/2021.03.27.437300

**Authors:** Carlos M. Duarte, David Ketcheson, Víctor M. Eguíluz, Susana Agustí, Juan Fernández-Gracia, Tahira Jamil, Elisa Laiolo, Takashi Gojobori, Intikhab Alam

**Affiliations:** Red Sea Research Centre (RSRC), King Abdullah University of Science and Technology, Thuwal 23955, Saudi Arabia; Computational Bioscience Research Centre (CBRC), King Abdullah University of Science and Technology, Thuwal 23955, Saudi Arabia; Computer, Electrical, and Mathematical Sciences and Engineering (CEMSE) Division, King Abdullah University of Science and Technology, Thuwal 23955, Saudi Arabia; Instituto de Física Interdisciplinar y Sistemas Complejos IFISC (UIB-CSIC), Palma de Mallorca, Spain

## Abstract

Evolutionary ecology theory provides an avenue to anticipate the future behavior of SARS-CoV-2. Here we quantify the accelerating evolution of SARS-CoV-2 by tracking the SARS-CoV-2 mutation globally, with a focus on the Receptor Binding Domain (RBD) of the spike protein believed to determine infectivity. We estimate that 384 million people were infected by March 1st, 2021, producing up to 10^21^ copies of the virus, with one new RBD variant appearing for every 600,000 human infections, resulting in approximately three new effective RBD variants produced daily. Doubling the number of RBD variants every 71.67 days followed by selection of the most infective variants challenges our defenses and calls for a shift to anticipatory, rather than reactive tactics.

**One-Sentence Summary:** Accelerating evolution of SARS-CoV-2 demands formulating universal vaccines and treatments based on big-data simulations of possible new variants.

## Main Text

Building on early ideas by Haldane (1), the evolutionary race between hosts and pathogens has been described, in a metaphoric sense, by the Red Queen theory(2). This metaphor refers to the warning of the Red Queen to Alice, in Lewis Carroll’s book (3), that in her kingdom *“it takes all the running you can do, to keep in the same place. If you want to get somewhere else, you must run at least twice as fast as that!”* (Carroll 1871).The repeated discovery of more infectious and possibly more deleterious variants of the SARS-CoV-2 virus shows that it is already engaged in this evolutionary race.

Here we provide insights into the current and future evolution of SARS-CoV-2 through a macroscopic consideration of evolutionary ecology theory. We first examine the rise of novel SARS-CoV-2 variants and their relationship to human infections, provide evidence of the rates and paths of selection driving the evolution of SARS-CoV-2 and, building on this evidence, discuss the expected outcomes and the most effective defense tactics. We focus our analysis on mutations at the 194 amino-acid RBD of SARS-CoV-2 (4) We do so on the basis of a unique resource, based on raw sequenced genomes from GISAID (www.gisaid.org) (5), identifying mutations and Mutation Fingerprints (MF), available through our in-house platform COVID-19 virus Mutation Tracker (CovMT; https://www.cbrc.kaust.edu.sa/covmt)(6). We define each set of SARS-CoV-2 genomes that generate the same amino acid sequence in the RBD region of the spike protein as a unique RBD variant (cf. methods).

### Virus production and mutation

The number of copies of the virus produced depends on the number of people infected globally along with the number of copies transcribed per infected subject. The verified number of diagnosed COVID-19 infections is an underestimate of the true infections, because it is constrained by the amount of testing. We, therefore, used a model based on the reported COVID-19 deaths (7) which is just over 2.6 million worldwide (as of March 1, 2021), based on demographics for each country along with age-specific infection fatality ratios (7). This model indicates that the true number of infections by March 1, 2021 exceeds 384 million (Fig 1a). Due to underreporting of COVID-19 deaths, the true total may be a few times above this estimate. Assuming a total transcription of about 10^9^ - 10^12^ viral genomes per individual along an infection cycle (8) and provided our cumulative estimate of infected persons, the number of SARS-CoV-2 copies produced globally since the pandemic started is in the range 10^17^ - 10^21^ as of March 1, 2021 (Fig. 1a).

**Figure 1.**
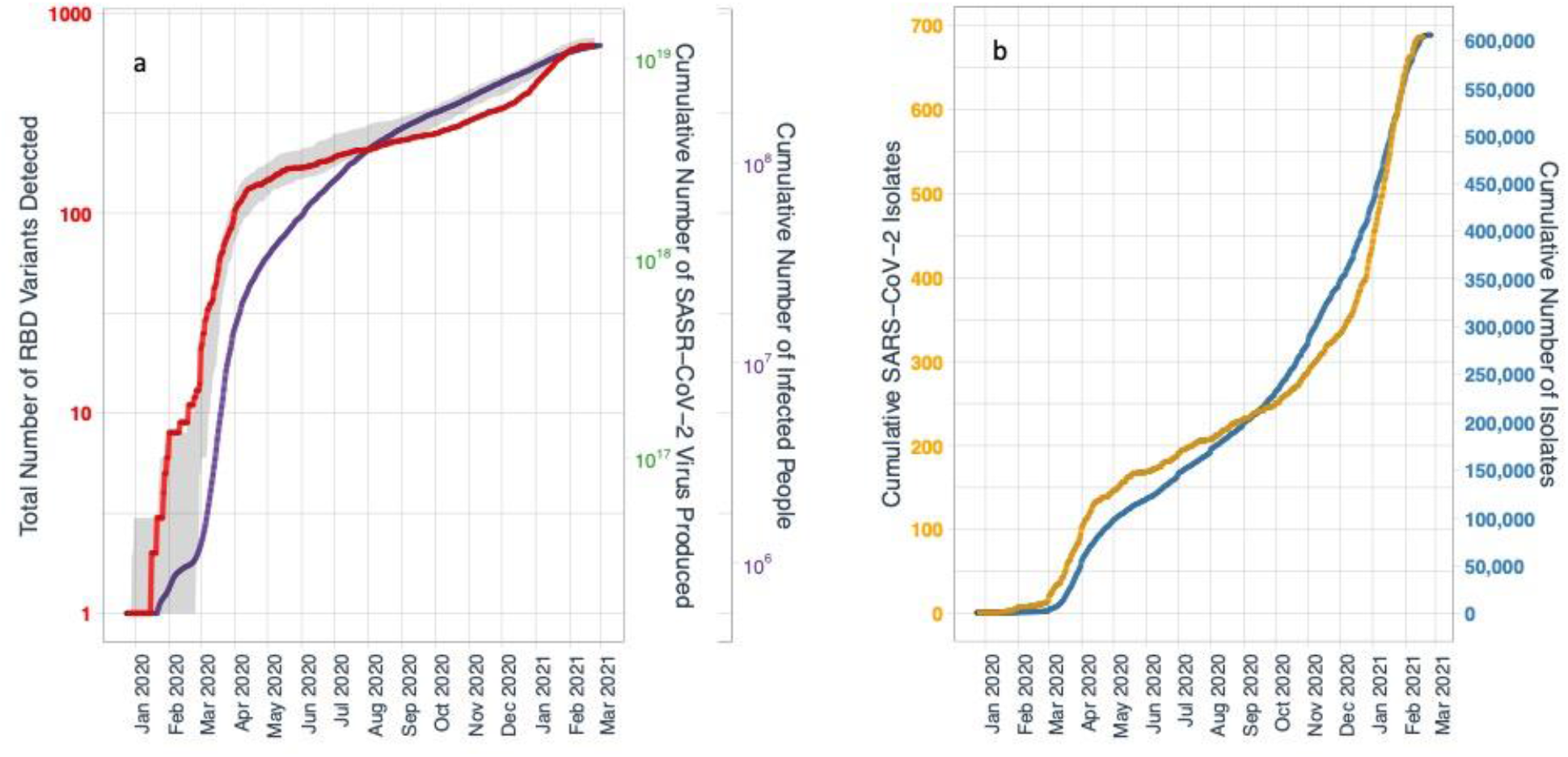
(a) The cumulative number of SARS-CoV-2-infected humans and calculated total number of virus copies produced until March 1, 2021 (cf. methods), and detected (red line) and predicted to be detected (grey shaded area, representing minimum and maximum bounds) RBD variants over time. The predicted RBD variants detected are based on Monte Carlo simulation of viral mutations, with a fixed rate of mutation per human infection and a Pareto distribution of variant fitness (see methods). Isolates are modeled as random samples from the resulting viral population. (b) Cumulative number of SARS-CoV-2 isolates sequenced and number of detected RBD variants over time. One new variant is reported for every 900 sequenced isolates.

The number of RBD variants rises in general correlation with the cumulative number of virus copies produced (Fig. 1a) but correlates much more strongly with the number of isolates sequenced (Fig. 1b), to reach 689 unique variants by March 1, 2020, consistent with mutation rate being a function of the number of copies produced (Fig. 1a), as every genome transcription is accompanied by a likelihood of mutations. The total number of RBD variants is growing with a doubling time of 71.67 ± 0.06 days (mean ± SE), accelerating from a maximum of about 237 days achieved at 291 days following the first detection in December 24, 2019 (Fig. 2a). The rapid rise in the number of variants at the onset of the pandemic (Fig. 1b) indicates that there was already significant diversity in December 2019, consistent with conclusions reported by the WHO mission (9). Many of the RBD variants present mutations elsewhere in their genome, so that each RBD variant, considering mutations in other parts of the genome, is represented by several variants, leading to a total of 90,744 unique variants presenting at least one mutation in the RBD region. Many sequenced isolates present mutations in regions of the genome other than the RBD domain, so the total number of SARS-CoV-2 variants (i.e. unique sequences across the entire genome) is much greater (320,762 as of March 1, 2021).

**Figure 2.**
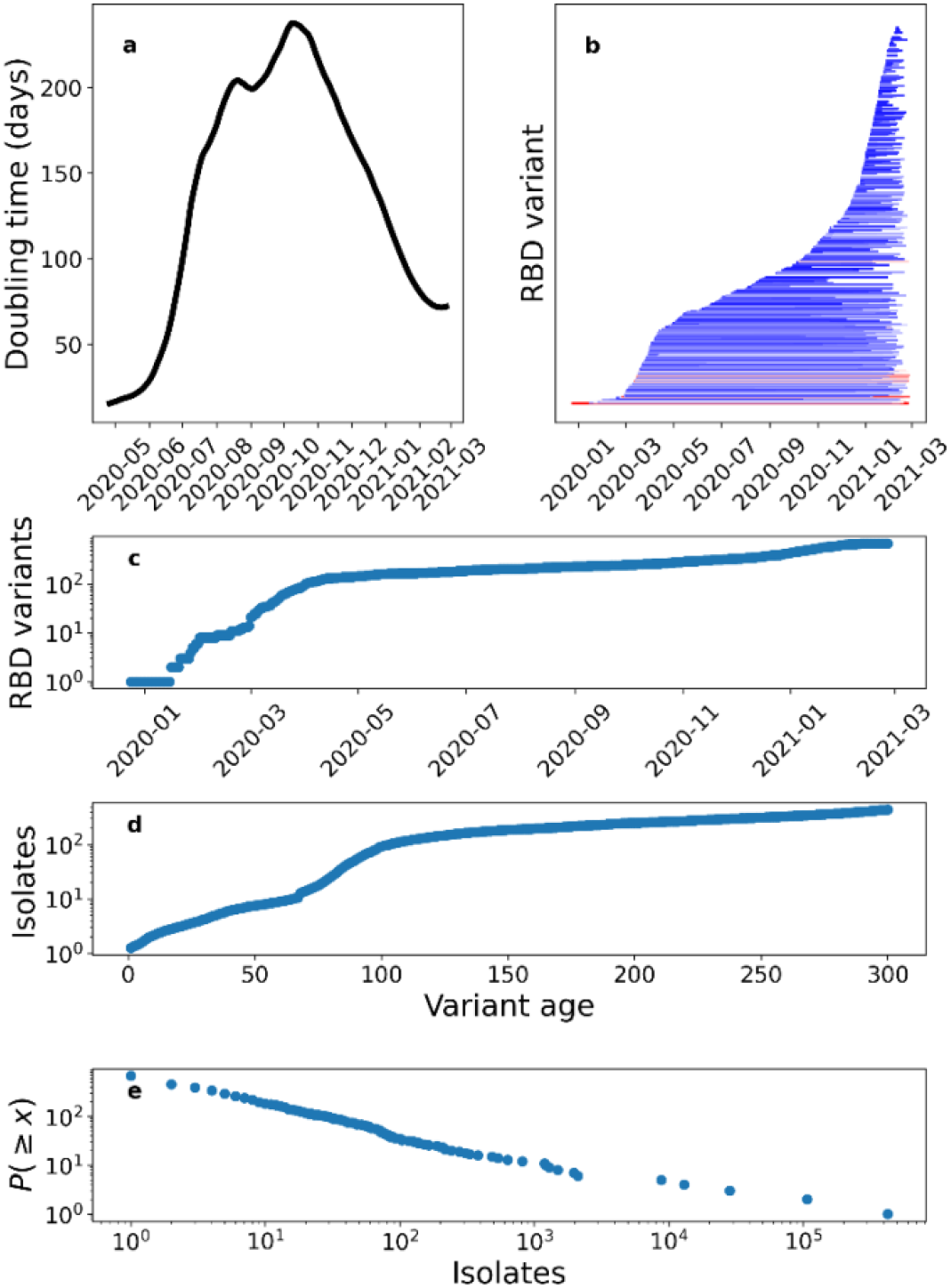
(a) Doubling time for detected RBD variants, calculated along a 100-day rolling window. (c) Rank abundance distribution of RBD variants. (b) The growth in the number of genome sequences of different unique RBD variants detected from SARS-CoV-2 isolates over time. The figure shows a line for each RBD variant along the y-axis that spans from the first day it was observed to the last date it was observed. The color scale denotes the final number of isolates of each RBD variant (increasing from blue to red). Yule’s model of the growth of RBD variants and isolates for each variant, showing the total number of detected unique RBD variants (c) and their age (as the time since their first detection) (d) over time, resulting in a power law rank abundance distribution (e). Numerically we find that the characteristic exponent of the exponential growth for RBD variants is 0.003532 ± 0.00003 day^-1^ (SD); then for the average growth, 0.00532 ± 0.00003 day^-1^ (SD); leading to an expected exponent of the rank abundance distribution (e) of 1.66 (to be compared with 1.60 from the data).

The number of unique RBD variants detected is certainly an underestimate of the number of variants in circulation, as their detection depends on sequencing effort, with a total of 616,389 reported genome sequences for SARS-CoV-2 isolates by March 1, 2021 (Fig. 1b). The rate of detection of new variants strongly depends on sequencing effort, with the ratio between these consistently around 1:900 (Fig. 1b). Direct modeling of the virus mutation is also capable of reproducing the observed data (Fig. 1a). Our model (cf. methods) suggests that the detected RBD variants represent about 85% of all variants, if we count only variants whose fitness leads to them comprising at least 0.0001% of the global viral population. This may be an optimistic estimate since real-world isolates represent a biased sample. The model suggests that one such new variant appears on average for every 600,000 human infections, or equivalently one new effective mutation per 10^16^ viral transcriptions. Given that our model estimated 2 million infections daily as of Feb. 15, this implies that three new effective variants (i.e. recruited to the virus population) are produced, on average, every day. Reproducing the initial rapid rise in the number of detected variants in March 2020 (Fig. 1b) requires a significant level of viral diversity in the initial pool, consistent with recent evidence of a larger-than-reported initial outbreak (9).

### SARS-CoV-2 evolution and selection

Evolutionary processes lead to genetic diversification along a branching process, with the evolutionary tree for SARS-CoV-2 RBD variants (Fig. 3a) characterized by a scaling between cumulative branching length and the 1.5 power of subtree size. The cumulative branching length is related to the mean subtree depth, depth = C/A, thus implying that the mean subtree depth scales as the square root of size (Fig. 3a). This scaling is characteristic of protein phylogenies, and deviates from fully balanced (i.e., resulting from lack of selection) and fully imbalanced trees (10). This evolutionary tree structure is consistent with non-random universal inferred patterns of evolution across scales, from the molecular level [e.g. protein families (10)] to phylogenetic differentiation ranging from micro-evolutionary to macro-evolutionary processes, shaping the diversity of life on the planet (11).

**Figure 3.**
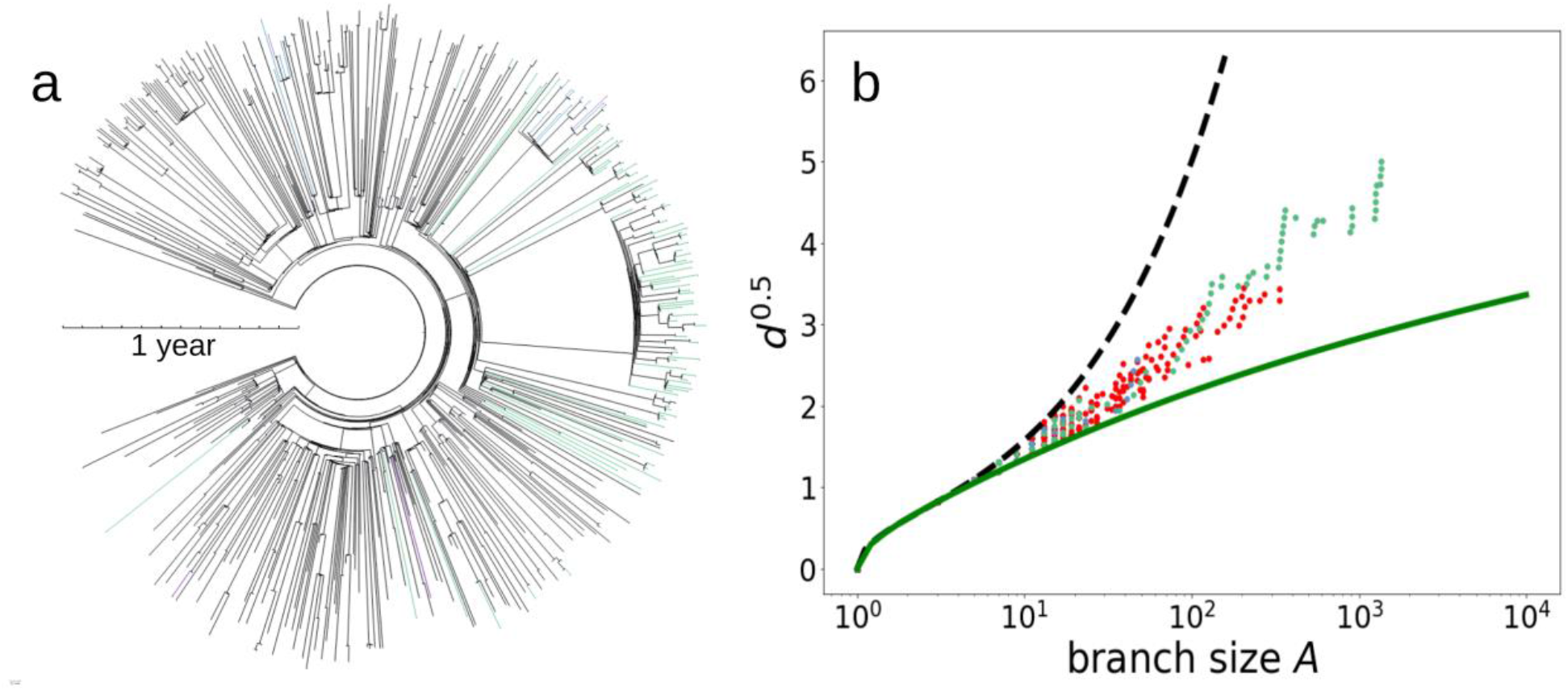
(a) Evolutionary tree of the 679 unique SARS-CoV-2 RBD variants detected by March 1, 2021. The so-called UK (B.1.1.7), South African (B.1.351) and Brazilian (P.1 as well as P.2) RBD variants are identified as green, blue and purple terminal leaves in the tree, respectively. The distance of the branching to the core represents, as indicated by the scale, the time when this branching occurred. (b) The relationship between the mean depth (d) and weight of the (A) cluster subtrees in the evolutionary tree. The discontinuous and continuous lines correspond to the two theoretical extreme binary trees: fully asymmetric trees (black dashed line) and fully symmetric (solid green line). The scales of the axes are chosen so that the behavior of the type d ~ (ln A)^2^ appears as a straight line Symbol colors correspond to the so-called UK (green), South African (blue) and Brazilian (purple) RBD variants.

Some recently identified RBD variants have rapidly risen to be as represented in the population of sequenced isolates as those identified many months before (Figs. 2c and 3a, Video S1, Fig. S1). These highly successful variants include, specifically, the so-called UK variant (B.1.1.7), in which a specific mutated amino acid sequence seems to have appeared independently multiple times (Fig. 3a). Indeed, the UK variant has diversified more and faster than other lineages in the RBD region (Fig. S1). As a result, the branch containing most of the UK variants has diversified greatly leading to heavy branches in the SARS-CoV-2 RBD evolutionary tree, with one of the branches rapidly diversifying over the past months (Fig. 3a, Video S1). The South African (B.1.351) and Brazilian (P.1 as well as P.2) RBD variants (Fig. 3), are also being clearly selected in the SARS-CoV-2 RBD evolutionary tree (Fig. 3, Video S1). A further indication of rapid evolution and selection is the rapid progression of the development of a hierarchical distribution of the abundance of the various RBD variants (Video S1), to conform to a power law consistent with the Yule law (Fig. 2c-e), a long-standing empirical observation for large groups of organisms (12) which requires exponential growth in the number of taxa (here RBD variants) in a lineage (here SARS-CoV-2) followed by exponential growth within each variant, as clearly seen in the evolutionary tree retrieved for SARS-CoV-2 RBD variants (Fig. 3a).

### Evolutionary outlook for the race between SARS-CoV-2 and its human hosts

There has been much discussion on the role of impacts on biodiversity in facilitating the arousal of zoonosis (13). However, the role of the huge, globally-connected human population, in the massive production of viruses propelling the rapid evolution of SARS-CoV-2 has not been sufficiently acknowledged.

Our analysis provides evidence for extraordinarily rapid evolution and selection of SARS-CoV-2, with the number of unique RBD variants currently doubling every 71.67 days, which has clearly reached a full speed in the Red Queen race, risking outpacing that of human defenses.

The same RBD variant, or identical-sequence variant, may arise independently in different locations, as the evolutionary tree suggests for the amino acid sequence shown by the UK variant (Fig. 3a). However, the SARS-CoV-2 evolutionary process deviates from random, with unbalanced branch development providing evidence of strong selection (Fig. 3a,b), consistent with the dynamics observed for the phylogenesis of protein families (10). Selection processes remove branches that are not infective while leading to heavy branches of the more infective strains (Fig. 3a). Indeed, the number of copies of the different RBD variants over time is not random, but are under selective pressure, particularly determined by the infectivity of the new variants emerged, as documented for the recently reported so-called UK (B.1.1.7), South African (B.1.351) and Brazilian (P.1 as well as P.2) RBD variants of SARS-CoV-2 (Video S1, Fig. S1). The result of this process is a highly hierarchical dynamic distribution of RBD variants, with a rank-abundance structure conforming to Yule’s law (12), implying that 1% of the variants contain more than 96% of the total isolates, with the 10 most detected variants by March 1, 2021, which include sequences of the UK and Brazilian variants, comprising 97.7 % (590750 out of 604572) of the total isolates sequenced (Fig. 2c-e). New, highly infective variants can rapidly recruit to this dominant pool (Video S1).

High mutation rates of RNA viruses, caused by error-prone RNA-dependent RNA polymerases (14) along with the huge virus production mediated by the huge pool of available human hosts propel the rapid evolution of SARS-CoV-2. The presence of a large number of variants in circulation within the same host population activates an additional mechanism, reassortment, for virus diversification. Reassortment involves the exchange of genetic segments between different strains of a segmented virus that have co-infected the same cell (14) which likely contributes to the rapid diversification of SARS-CoV-2. Provided a doubling time of SARS-CoV-2 RBD variants of 71.67 days, the number of SARS-CoV-2 RBD variants will continue to expand. This rapid diversification and selection of RBD variants characterize the evolutionary “Red Queen” race and predicts the selection of more infective variants becoming dominant in a highly hierarchical distribution dynamically conforming to Yule’s law. This heralds a new phase in the pandemic characterized by accelerating evolutionary rates of the virus, which will impose new challenges as new variants of concern add to those already detected in the UK, South Africa and Brazil.

Mutation and, possibly, reassortment propel SARS-CoV-2 to be running fast in terms of the Red Queen Theory. As the Red Queen advised Alice, we need to run even faster, just to keep in place, and much faster if we are to overcome the pandemic well before this declines upon reaching the limitation of available hosts. The Red Queen theory posits that hosts develop evolutionary defenses through recombination under sexual reproduction allowing them to modify their genome to anticipate and prevent pathogen attacks (2, 15, 16) This requires selection across generations and catastrophic mortality for SARS-CoV-2 morbidity to be selected against. Our defense mechanisms include protections to avoid contact with the virus, and therapies and vaccines once SARS-CoV-2 enters our bodies. External defenses include social distancing, with strict lockdowns proven across many nations to be the most effective defense mechanism, whatever unpopular, to contain the pandemic, along with wearing protections and emerging uses of nanotechnology for virus detection and interception (17, 18) This effort must be complemented with the continuous development of a diverse suite of universal therapies, such as multivalent nanobodies (19) and vaccines, eliciting immune defenses that vary and can defend us against a wide range of RBD variants, existing and forthcoming, as new variants that overcome immune defenses produced by previously infected or vaccinated people arise, as demonstrated by our long experience in coping with the drift and shift of the influenza virus (20). Indeed, recent reports indicate that the convalescent sera and BNT162b2 mRNA vaccine may not be effective, or may be only partially so, against the S. African B.1.351 variant (21).

Evolutionary ecology theory helps formulate predictions on the future behavior of SARS-CoV-2. On the other hand, the COVID-19 pandemic provides an unprecedented opportunity to test evolutionary ecology theory, which has been largely inferential in nature. This is important as never before had an evolutionary process been tracked in real time and with such wealth of openly available genomic data at a global scale. The SARS-CoV-2 validates a number of evolutionary theories and laws, such as the evolutionary underpinning of the partially imbalanced architecture of phylogenetic trees across evolutionary scales (10, 11) the diversification process responsible for the long-standing Yule law (12), and the more targeted framework of the Red Queen theory (2) predicting the evolutionary tactics of pathogens.

The development of the vaccine in record time, a feat rendered possible by unprecedented collaboration, has been celebrated as the start of the end of the pandemic. Rather, it may be the beginning of a new phase, where the continuous development of novel and diverse and universal vaccines, which requires sustained global collaboration, will be, along with effective human confinement, our best tools to avoid the rapid evolution of SARS-CoV-2 from outpacing us in the race to end the pandemic. *In silico* analysis of the effectiveness of current vaccines against plausible RBD variants not yet detected, and the design of new effective vaccines against such variants will enable us to overtake SARS-CoV-2 in the evolutionary race, as a reactive, catch-up tactic, as that played to date, will carry continuous risks. Artificial Intelligence may further help analyze the immunogenicity of all the nonsynonymous variations across described and predicted SARS-CoV-2 sequences to generate a blueprint for effective vaccine development (22, 23).

Unprecedented in the history of human pandemics, we now have the massive, global genomic data, supercomputers to analyze them and understanding of evolutionary processes required to predict and anticipate future possible evolutionary pathways for SARS-CoV-2. Extreme computing, enabled by collaborative sharing of genomic data, may arise as the new weapon in our “Red Queen” race against human pathogens.

## Acknowledgments

This research was funded by King Abdullah University of Science and technology through research made available to the Computational BioScience Research Center, CMD and TG.

## Author contributions

Conceptualization: CMD and SA

Methodology: IA, DK, VME, JFG

Investigation: all authors

Visualization: TJ, JFG, VME

Funding acquisition: CMD, TG

Project administration: CMD, IA

Supervision: CMD, TG

Writing – original draft: CMD, SA, DK, EL, JFG, IA

Writing – review & editing: all authors

## Data and materials availability

Mutation Fingerprints (MF) derived classification of SARS-CoV-2 isolates into MF variants grouped by sampling dates and locations, updated daily, is available on the download section of CovMT website, https://www.cbrc.kaust.edu.sa/covmt. Supplementary Materials

## Supplementary Materials

### Materials and Methods

The number of positive tests for COVID-19 virus infections reached ~100 million in January, 2021; see https://covid19.who.int. To infer real infections over time, we use numbers of confirmed deaths based (24, 25). We apply a regularized deconvolution with an estimated time-to-death distribution to infer the number of real infections over time, as described in Ketcheson *et al*. (7).

#### COVID-19 virus Isolate Genomes

Largest resource of Isolate genomes in COVID-19 virus is available at the **G**lobal **I**nitiative on **S**haring **A**vian **I**nfluenza **D**ata (GISAID, www.gisaid.org)(5). As of March 01, 2021, more than 616,389 SARS-CoV-2 genomes are available from around the world.

#### COVID-19 virus variants

Mutations in the genome of SARS-CoV-2 are the basis to define its genomic variants. There are several ways to group mutations in COVID-19 virus. GISAID provides generic clades, and more detailed lineages are provided by Phylogenetic Assignment of Named Global Outbreak LINeages (PANGOLIN) tool by Rambaut *et al*.(26, 27)

In our effort of a daily updated COVID-19 virus Mutation Tracking system (CovMT, https://www.cbrc.kaust.edu.sa/covmt)(6), we provide mutation fingerprints (MFs). A mutation fingerprint is defined based on all synonymous and nonsynonymous mutations in an isolate genome. To avoid noise and sequencing errors, a minimum frequency of a mutation from the global population of isolate genomes is kept at 0.001%. We include information about GISAID clades and PANGOLIN lineages for easy exploration of variants. A daily updated table on counts of MFs grouped by sampling dates and location is available at https://www.cbrc.kaust.edu.sa/covmt/data/Variants/World/World_variants_summary.zip

#### RBD variants

The Receptor Binding Domain (RBD) region of Spike protein in SARS-CoV-2 is an important domain region that facilitates the binding of this virus to host cells. Unique RBD variants are defined as those showing exactly the same amino acid sequence for the RBD region of the Spike protein for SARS-CoV-2. We group SARS-CoV-2 genomes into RBD variants by taking the subset of Mutation Fingerprints restricted to the RBD region only and considering only the amino acid mutations. To avoid sequencing errors only those mutations are considered where mutation frequency at the population level is 0.001%, an error rate defined based on Illumina base call quality score of Q25-30. Majority of the SARS-CoV-2 genomes submitted to GISAID as based on Illumina sequencing technology. Mutation Fingerprints of RBD variants as well as all other variants with associated metadata are available at CovMT webpage, https://www.cbrc.kaust.edu.sa/covmt/index.php?p=world-variants.

#### Mutation modeling

We estimate the number of effective mutations per human infection based on a direct modeling approach using Monte Carlo simulation. New RBD variants are generated based on an assumed mutation rate (chosen to fit the data). The fitness of new variants is taken as a Pareto distribution with scale 10^-6^ and shape parameter ¼. The proportional population of two variants is assumed to change at a rate proportional to the ratio of their fitnesses, with a characteristic time of twenty days. Isolates are modeled as random samples from the resulting viral population.

#### Phylogenetic tree for loose RBD variants

We use the phylogenetic tree provided by nextstrain (https://nextstrain.org/ncov/global accessed 11/02/2021), which provides the phylogeny of ~4000 genomes sampled between december 2019 and February 2021. We identified which samples had any mutation in the RBD region and pruned the tree to contain only those. We end up with 900 different RBD loose variants. We also identified which of these variants belonged to the UK (B.1.1.7), South African (B.1.351) and Brazilian (P.1 and P.2) variants using data provided by COVID-19 virus mutation tracker (https://www.cbrc.kaust.edu.sa/covmt/). For the analysis of the depth scaling of the tree branches we followed the procedure described in refs.

#### Fit to power laws

The fits to power laws in Fig. 2 were performed using maximum likelihood (28).

**Fig S1.**
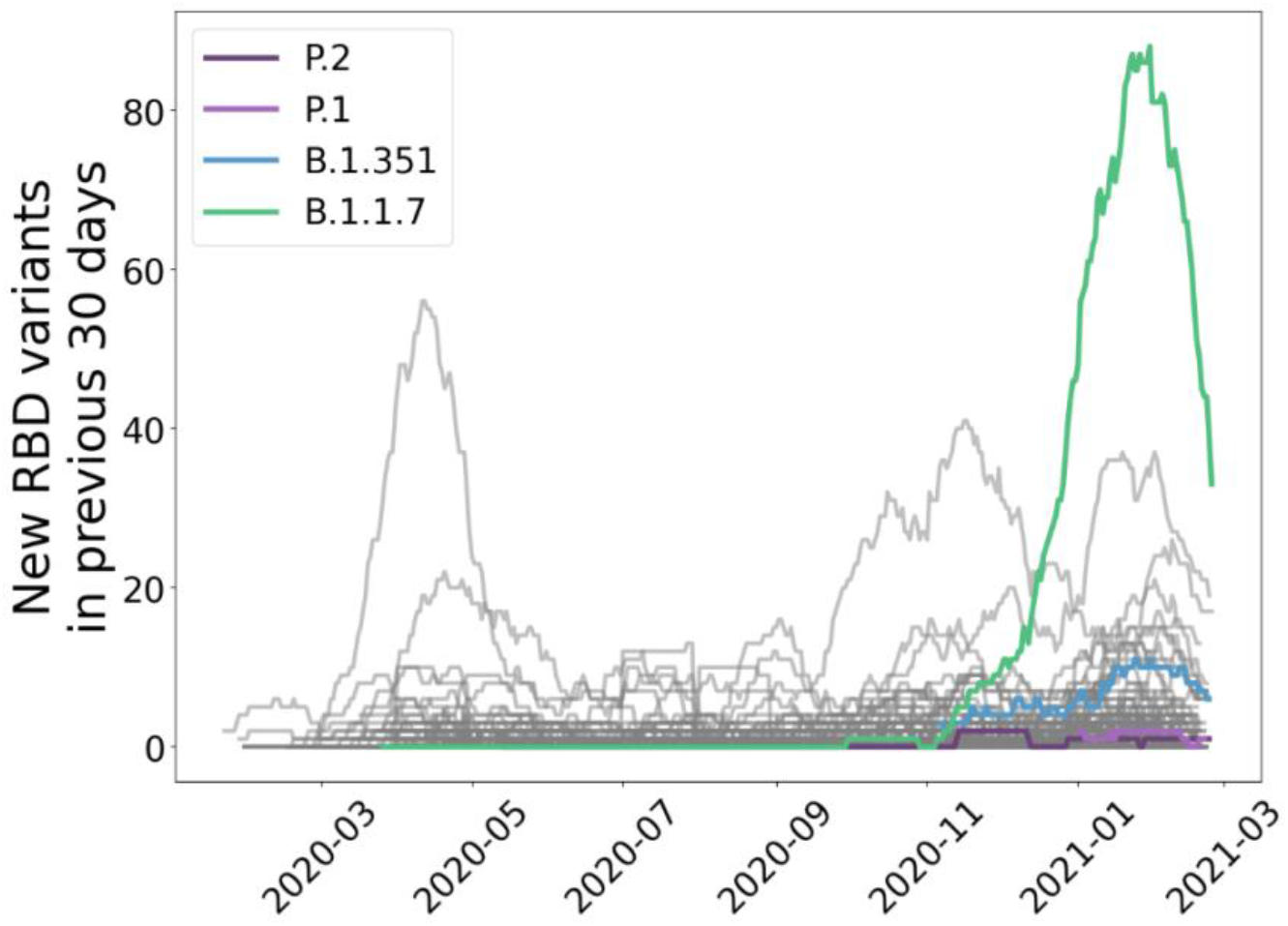
The rate of discovery of new unique RBD variants in the preceding 30 days for each lineage in the SARS-CoV-2 evolutionary tree (Fig. 3a). We identify the UK lineage (B.1.1.7) in green, the South African lineage (B.1.351) in blue and the Brazilian lineages (P.1 and P.2) in different shades of purple. In grey are all the other lineages. While the South African and Brazilian variants do not stand out, the UK variant shows a clear excess in diversification in the RBD region as compared with the other lineages, with a peak of more than 80 new variants observed in the previous month at the end of January.

### Movie S1

An animation of the rank distribution of SARS-CoV-2 RBD variants over time, showing the dynamic replacement of new variants among the top 10 variants, in terms of their total representation in the pool of sequenced isolates. As also shown in Figure 2, by mid-February 2020, the SARS-CoV-2 population is dominated by the original RBD variant and the RBD:1(N501Y;A23063T) variant. SARS-CoV-2 variants with N501Y mutation were first detected in Brazil in April 2020. These variants re-appeared in the UK with more mutations outside the RBD domain during September 2020 and are referred to as the “UK variant” by the media (i.e. B.1.1.7 lineage). The second dominant variant by February 2020 is the RBD:1(S477N;G22992A), first identified in March 2020 in Australia and France, becoming dominant in Australia during summer with spread to several other countries to remain dominant until December 2020, when the “UK variant” exceeded its abundance. The third variant most found in isolates by mid-February 2021 is the S_RBD:1(N439K;C22879A), first detected in the UK in March 2020, increasing in number of isolates sequenced after October 2020 in the UK and Denmark. Another RBD variant with an increase in the number of SARS-CoV-2 genomes sequenced, just after L452R (dominant in the United States of America), is the one with mutation E484K. Variants with E484K mutation got attention when appeared with three RBD mutations RBD:3(K417N;G22813T|E484K;G23012A|N501Y;A23063T) referred to as the “South African variant”, lineage B.1.351. SARS-CoV-2 variants with E484K mutation are able to evade antibodies with reduced efficacy from existing vaccines21,30,31. Furthermore, the highly infectious so-called UK variant is now confirmed to have acquired the mutation E484K2 it is expected that the number of isolates with mutation E484K will increase in the future. A timeline of the top 10 RBD variants can be seen in Figure S1 and at CovMT dedicated page, https://www.cbrc.kaust.edu.sa/covmt/index.php?p=top-rbd-variants-line

## References and Notes

1. J. B. S. Haldane, Disease and evolution. Supplement to La Ricerca Scientifica 19, 68–76 (1949).

2. G. Bell, The Masterpiece of Nature: the Evolution and Genetics of Sexuality. (Univ. Calif. Press, 1982).

3. L. Carroll, Through the looking-glass, and what Alice found there. (Macmillan, 1871).

4. J. Lan, J. Ge, J. Yu, S. Shan, H. Zhou, S. Fan, Q. Zhang, X. Shi, Q. Wang, L. Zhang, X. Wang, Structure of the SARS-CoV-2 spike receptor-binding domain bound to the ACE2 receptor. Nature, 581, 215–220 (2020).

5. S. Elbe, G. Buckland-Merrett, Data, disease and diplomacy: GISAID’s innovative contribution to global health. Global Challenges, 1, 33–46 (2017).

6. I. Alam, A. Radovanovic, R. Incitti, A. A. Kamau, M. Alarawi, E. I. Azhar, T. I. Gojobori, CovMT: an interactive SARS-CoV-2 mutation tracker, with a focus on critical variants. Lancet Infect. Dis. (2021).

7. D. I. Ketcheson, H. C. Ombao, P. Moraga, T. Ballal, C. M. Duarte, http://medrxiv.org/lookup/doi/10.1101/2020.05.11.20097972 (2020).

8. R. Sender, Y. M. Bar-On, A. Flamholz, S. Gleizer, B. Bernsthein, R. Phillips, R. Milo, https://www.medrxiv.org/content/10.1101/2020.11.16.20232009v1 (2020).

9. N. Paton Walsh, “CNN Exclusive: WHO Wuhan mission finds possible signs of wider original outbreak in 2019”. CNN, retrieved 15-02-2021. https://edition.cnn.com/2021/02/14/health/who-mission-china-intl/index.html

10. A. Herrada, V. M. Eguíluz, E. Hernández-García, C. M. Duarte, Scaling properties of protein family phylogenies. BMC Evol Biol 11, 155 (2011).

11. A. Herrada, C. J. Tessone, K. Klemm, V. M. Eguíluz, E. Hernández-García, C. M. Duarte, Universal Scaling in the Branching of the Tree of Life. PLoS ONE 3, e2757 (2008).

12. J. C. Willis, G. U. Yule, Some statistics of evolution and geographical distribution in plants and animals, and their significance. Nature, 109, 177 (1922).

13. J. Tollefson, Why deforestation and extinctions make pandemics more likely. Nature, 584, 175–176 (2020).

14. B. D. Greenbaum, E. Ghedin, Viral evolution: beyond drift and shift. Current opinion in microbiology, 26, 109–115 (2015)

15. K. Clay, P. X. Kover, The Red Queen hypothesis and plant/pathogen interactions. Annual Review of Phytopathology, 34, 29–50 (1996)

16. M. Salathé, R. D. Kouyos, S. Bonhoeffer, The state of affairs in the kingdom of the Red Queen. Trends in Ecology & Evolution, 23, 439–445 (2008)

17. M. Srivastava, N. Srivastava, P. K. Mishra, B. D. Malhotra, Prospects of nanomaterials-enabled biosensors for COVID-19 detection. Science of The Total Environment, 754, 142363 (2021).

18. A. D. Chintagunta et al, Nanotechnology: an emerging approach to combat COVID-19. Emergent Materials, 1–12 (2021).

19. P. A. Koenig, H. Das, H. Liu, B. M. Kümmerer, F. N. Gohr, L. M. Jenster, L. D. J. Schiffelers, Y. M. Tesfamariam, M. Uchima, J. D. Wuerth, K. Gatterdam, N. Ruetalo, M. H. Christensen, C. I. Fandrey, S. Normann, J. M. P. Tödtmann, S. Pritzl, L. Hanke, J. Boos, M. Yuan, X. Zhu, J. L. Schmid-Burgk, H. Kato, M. Schindler, I. A. Wilson, M. Geyer, K. U. Ludwig, B. M. Hällberg, N. C. Wu, F. I. Schmidt, Structure-guided multivalent nanobodies block SARS-CoV-2 infection and suppress mutational escape. Science, 371 (2021).

20. H. Kim, R. G. Webster, R. J. Webby, Influenza virus: dealing with a drifting and shifting pathogen. Viral immunology, 31, 174–183 (2018).

21. T. Tada, B. M. Dcosta, M. Samanovic-Golden, R. S. Herati, A. Cornelius, M. J. Mulligan, N. R. Landau, https://www.biorxiv.org/content/10.1101/2021.02.05.430003v1 (2021)

22. T. N. Starr, A. J. Greaney, S. K. Hilton, D. Ellis, K. H. D. Crawford, A. D. Dingens, M. J. Navarro, J. E. Bowen, M. A. Tortorici, A. C. Walls, N. P. King, D. Vessler, J. D. Bloom, Deep mutational scanning of SARS-CoV-2 receptor binding domain reveals constraints on folding and ACE2 binding. Cell, 182, 1295–1310 (2020).

23. B. Malone, B. Simovski, C. Moliné, J. Cheng, M. Gheorghe, H. Fontenelle, I. Vardaxis, S. Tennøe, J. A. Malmberg, R. Stratford, T. C. Malone, Artificial intelligence predicts the immunogenic landscape of SARS-CoV-2 leading to universal blueprints for vaccine designs. Scientific reports, 10, 1–14 (2020)

24. COVID-19 Data Repository, Center for Systems Science and Engineering (CSSE), Johns Hopkins University, https://github.com/CSSEGISandData/COVID-19.

25. Coronavirus (Covid-19) Data in the United States, The New York Times, Retrieved 12-02-2021 https://github.com/nytimes/covid-19-data.

26. GISAID - Clade and lineage nomenclature aids in genomic epidemiology studies of active hCoV-19 viruses. https://www.gisaid.org/references/statements-clarifications/clade-and-lineage-nomenclature-aids-in-genomic-epidemiology-of-active-hcov-19-viruses/

27. A. Rambaut, E. C. Holmes, Á. O’Toole, V. Hill, J. T. McCrone, C. Ruis, L. du Plessis, O. G. Pybus, A dynamic nomenclature proposal for SARS-CoV-2 lineages to assist genomic epidemiology. Nat Microbiol 5, 1403–1407 (2020).

28. A. Clauset, C. R. Shalizi, M. E. J. Newman, Power-Law Distributions in Empirical Data. SIAM Rev., 51, 661–703 (2009).

29. D. A. Collier, A. De Marco, I. A. T. M. Ferreira, B. Meng, R. Datir, A. C. Walls, S. A. Kemp S, J. Bassi, D. Pinto, C. Silacci Fregni, S. Bianchi, M. A. Tortorici, J. Bowen, K. Culap, S. Jaconi, E. Cameroni, G. Snell, M. S. Pizzuto, A. Franzetti Pellanda, C. Garzoni, A. Riva, The CITIID-NIHR BioResource COVID-19 Collaboration, A. Elmer, N. Kingston, B. Graves, L. E. McCoy, K. G. C. Smith, J. R. Bradley, L. Ceron-Gutierrez, G. Barcenas-Morales, The COVID-19 Genomics UK (COG-UK) consortium, W. Harvey, H. W. Virgin, A. Lanzavecchia, L. Piccoli, R. Doffinger, M. Wills, D. Veesler, D. Corti, R. K. Gupta, D. A. Collier, https://www.medrxiv.org/content/10.1101/2021.01.19.21249840v4.full (2021).

30. H. Tegally, E. Wilkinson, M. Giovanetti, A. Iranzadeh, V. Fonseca, J. Giandhari, D. Doolabh, S. Pillay, E. J. S. N. Msomi, K. Mlisana, A. von Gottberg, S. Walaza, M. Allam, A. Ismail, T. Mohale, A. J. Glass, S. Engelbrecht, G. Van Zyl, W. Preiser, F. Petruccione, A. Sigal, D. Hardie, G. Marais, M. Hsiao, S. Korsman, M. A. Davies, L. Tyers, I. Mudau, D. York, C. Maslo, D. Goedhals, S. Abrahams, O. Laguda-Akingba, A. Alisoltani-Dehkordi, A. Godzik, C. Kurt Wibmer, B. Trevor Sewell, J. Lourenço, L. C. J. Alcantara, S. L. Kosakovsky Pond, S. Weaver, D. Martin, R. J. Lessells, J. N. Bhiman, C. Williamson, T. de Oliveira, https://www.medrxiv.org/content/10.1101/2020.12.21.20248640v1.full (2020).

